## Supplementary material for "Promiscuous structural cross-compatibilities between major shell components of *Klebsiella pneumoniae* bacterial microcompartments": Supplementary Figures and Tables.pdf

**A**

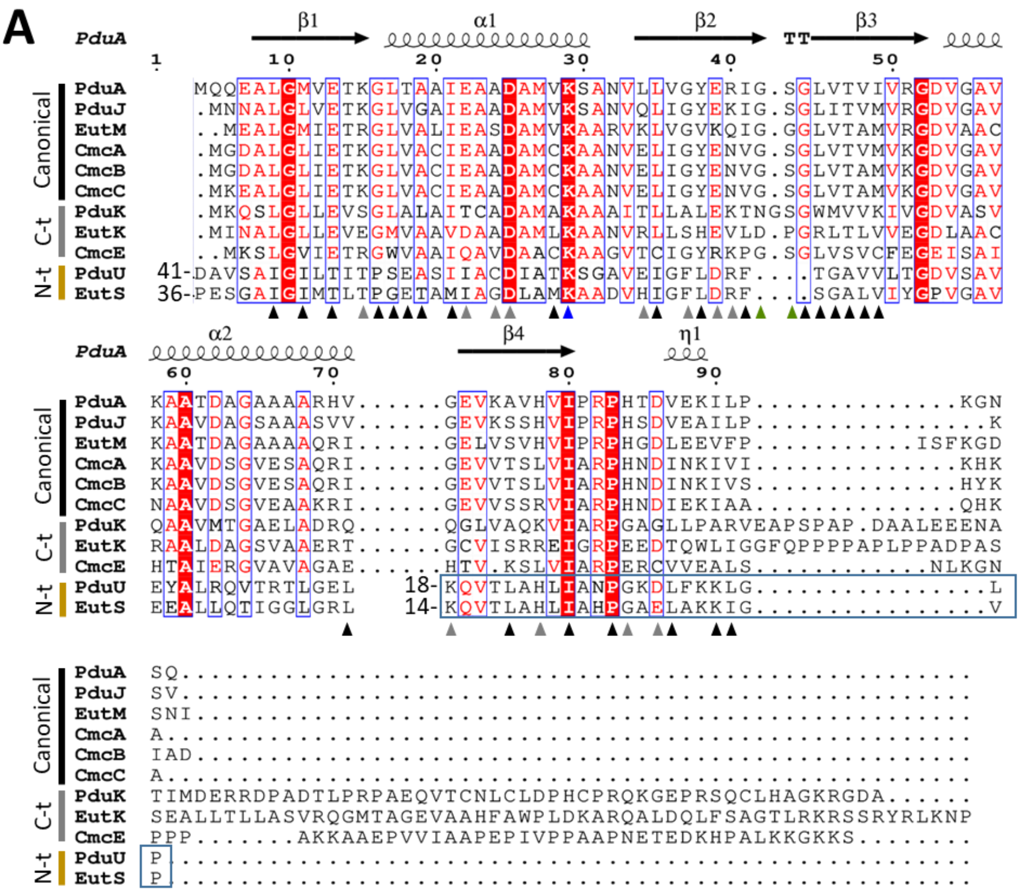

**B**

| | GRM2 | | | | EUT1 | | | PDU1A | | | |
|---|---|---|---|---|---|---|---|---|---|---|---|
| | CmcA | CmcB | CmcC | CmcE | EutK | EutM | EutS | PduA | PduJ | PduK | PduU |
| CmcA | | 95 | 86 | 56 | 42 | 56 | 22 | 63 | 66 | 41 | 26 |
| CmcB | | | 86 | 44 | 40 | 57 | 22 | 59 | 64 | 38 | 24 |
| CmcC | | | | 42 | 44 | 59 | 22 | 61 | 66 | 40 | 24 |
| CmcE | | | | | 27 | 38 | 21 | 40 | 45 | 32 | 20 |
| EutK | | | | | | 42 | 21 | 40 | 45 | 32 | 20 |
| EutM | | | | | | | 19 | 63 | 61 | 34 | 22 |
| EutS | | | | | | | | 20 | 22 | 18 | 56 |
| PduA | | | | | | | | | 77 | 35 | 25 |
| PduJ | | | | | | | | | | 39 | 23 |
| PduK | | | | | | | | | | | 20 |
| PduU | | | | | | | | | | | |

| GRM2 | | | | EUT1 | | | PDU1A | | | | |
|---|---|---|---|---|---|---|---|---|---|---|---|
| CmcA | CmcB | CmcC | CmcE | EutK | EutM | EutS | PduA | PduJ | PduK | PduU | |
| | 98 | 90 | 50 | 43 | 60 | 10 | 70 | 73 | 39 | 23 | GRM2 |
| | | 93 | 47 | 43 | 63 | 10 | 67 | 70 | 37 | 20 | |
| | | | 45 | 47 | 65 | 10 | 67 | 72 | 37 | 20 | |
| | | | | 27 | 40 | 10 | 47 | 48 | 34 | 20 | |
| | | | | | 38 | 18 | 37 | 43 | 29 | 18 | EUT1 |
| | | | | | | 15 | 63 | 63 | 29 | 23 | |
| | | | | | | | 15 | 12 | 17 | 73 | |
| | | | | | | | | 80 | 39 | 27 | PDU1A |
| | | | | | | | | | 41 | 23 | |
| | | | | | | | | | | 22 | |

**Figure S1. Comparison of *Kpe* BMC-H sequences.**

**A.** Sequence alignments were prepared with the RCSB pairwise alignment tool (TM-align method), taking as input the structures from the BMC-H monomers generated by AF2. Secondary structure elements from PduA are indicated as reference. EutS and PduU present secondary structure permutations. They were aligned to PduA using the JCE-CP (flexible) method and manually verified. The permutation is highlighted by the black rectangle, indicating the number of the first shown residue. PduU and EutS residues that build the N-terminal β-barrel were excluded. Residues fully or partly embedded at the interface between monomers are indicated by black or grey arrows, respectively. Also in part participating to the interface are critical lysine (K26 in PduA) and pore residues, identified by blue and green arrows, respectively. The presentation was generated online with ESPript 3.0. **B.** Percentage of sequence identity between *Kpe* BMC-H. Values based on either residues belonging to the common BMC-H core domain (left) or only those falling at the interface between monomers (right). Colored values are to highlight cases exhibiting more than 10% discrepancy of identity when comparing the two sets of residues.

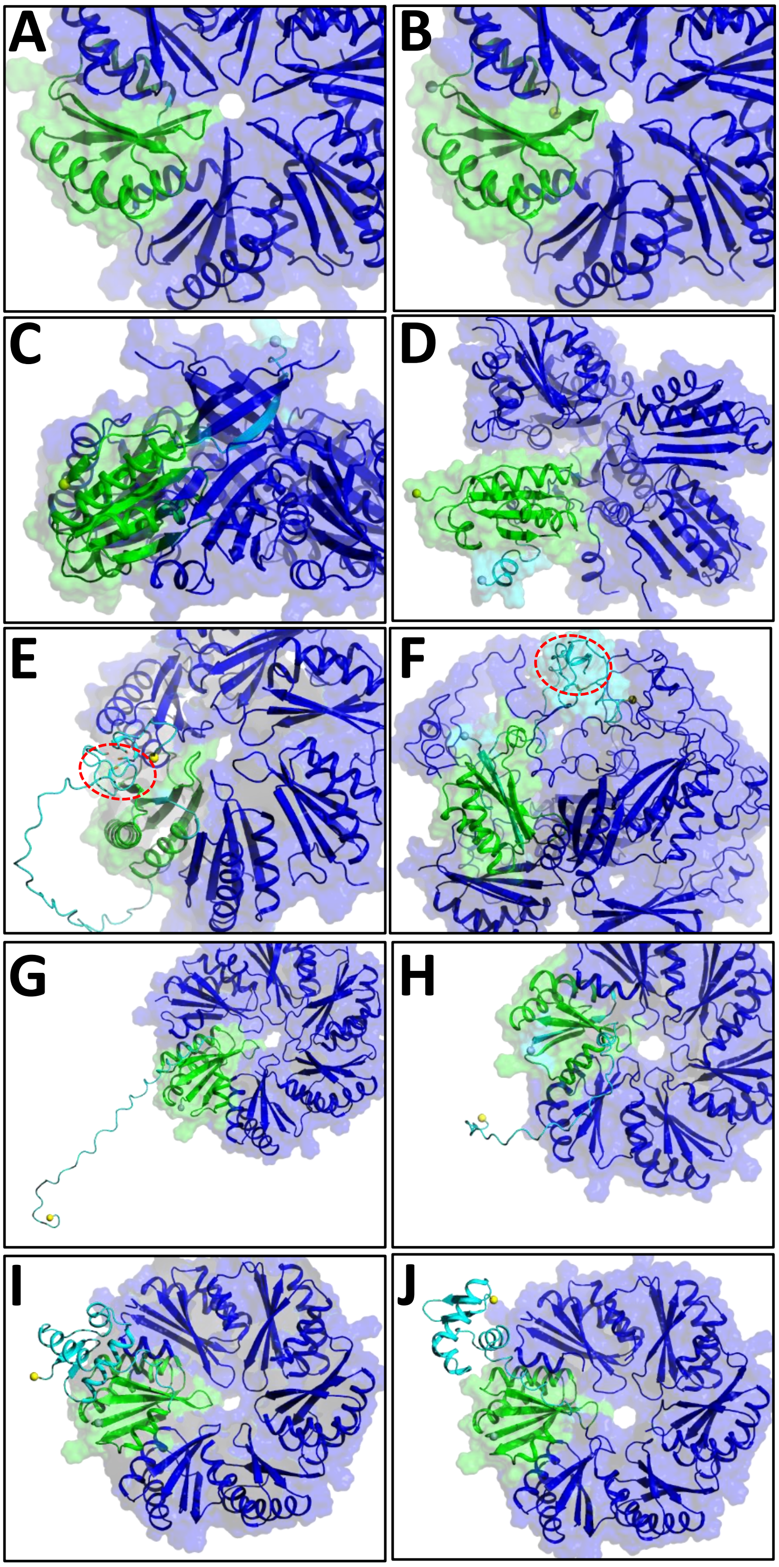

**Figure S2. Structural details on predicted BMC-H homo-hexamer models.**
Cartoon illustration of portions of representative BMC-H structures predicted by AF2 (left panels) or ESMFold (right panels). From top to bottom are presented the next BMC-H: CmcA (panels ***A,B***), EutS (***C,D***), PduK (***E,F***), CmcE (***G,H***) and EutK (***I,J***). The first monomer of the hexamer is colored green, all other monomers in blue. Views are from the hexamer convex side for CmcA and PduK, concave side for CmcE and EutK. In the case of EutS, a side view is shown. Please note that ESMFold did not predict hexamer for EutS (***D***) and PduK (***F***). The cysteine-rich domain of PduK is highlighted by the dashed red circles, with S-atoms appearing as orange sticks. To simplify the views, C-terminal extensions in the last three rows are shown only for the first hexamer (cyan). Light green and yellow spheres indicate the localization of the N- or C-terminal residue of the first monomer, respectively.

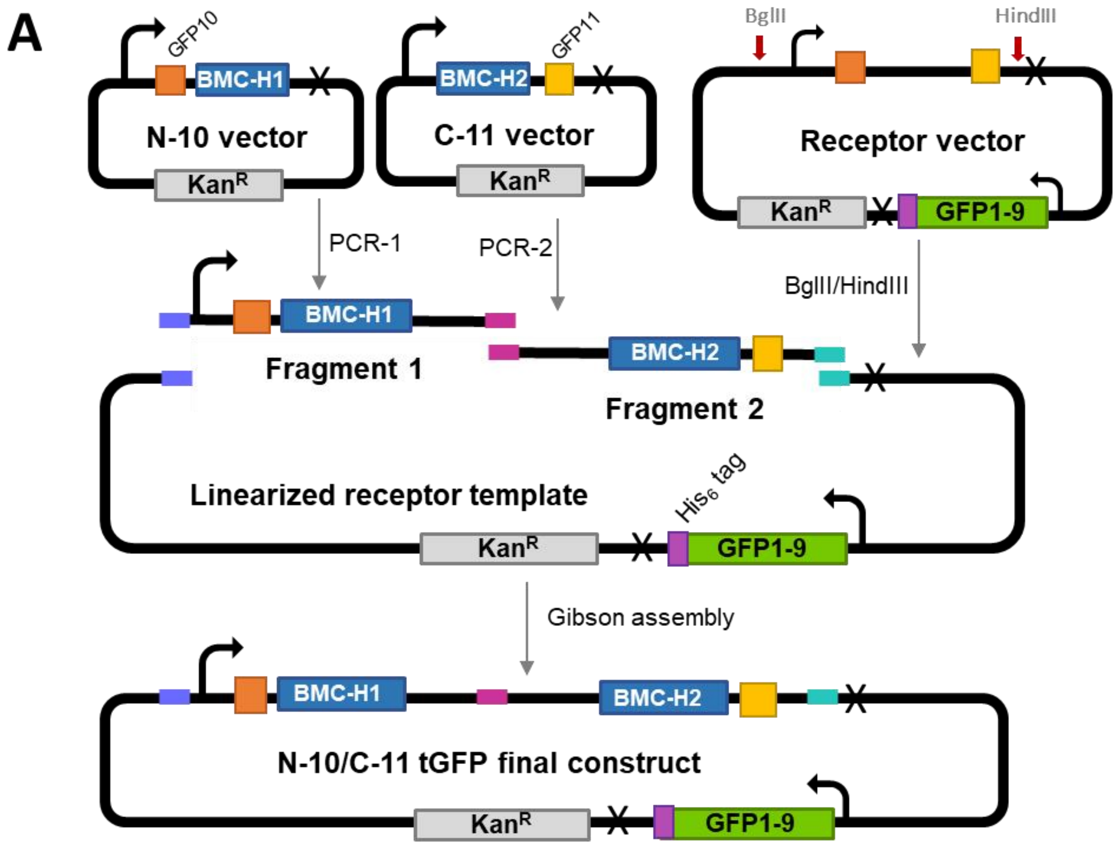

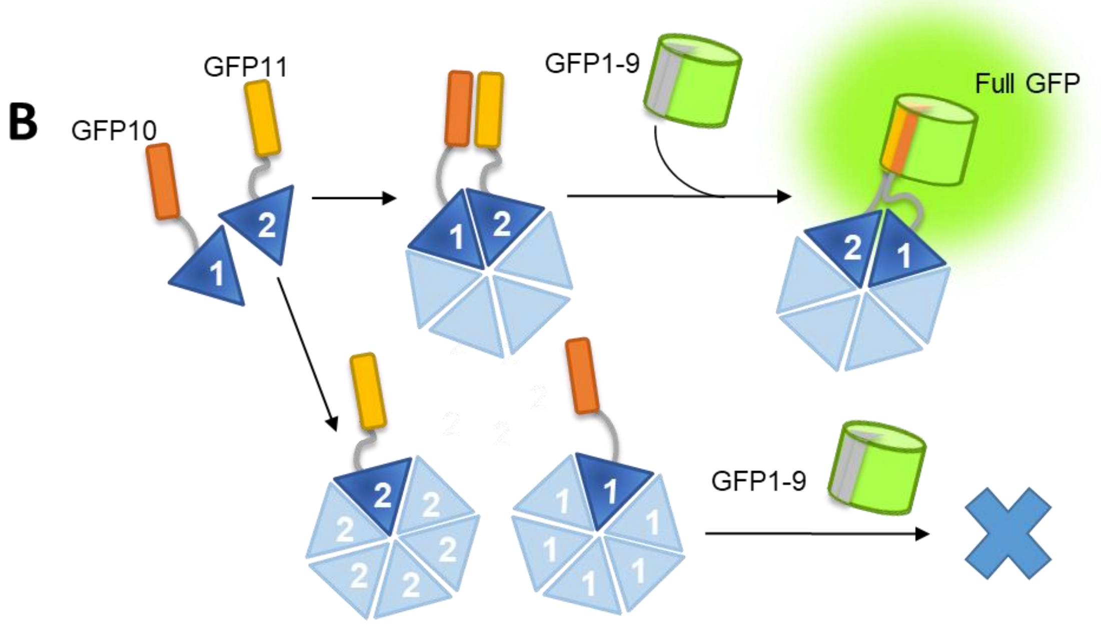

**Figure S3. The tripartite GFP screening approach.**

**A**. General strategy of construction of plasmids for tGFP assays. Preliminarly, plasmids coding for individual BMC-H with GFP10 or GFP11 tags in either C- or N-terminal were mounted on a pET26b-based template, giving rise to N-10, C-10, N-11 or C-11 vectors. Sequences coding for the two BMC-H of interest were then amplified by PCR from the corresponding plasmids. For simplicity, only one (N-10/C-11) of the eight possible combinations for a given pair of BMC-H is shown here. Primers included 15nt regions allowing hybridization to either the adjacent fragment (pink box) or to the receptor vector (blue and magenta). The tGFP receptor vector (Kan$^R$) included necessary information for the independent expression of the GFP1-9. The fragments and opened vector were Gibson-assembled giving rise to the final tGFP construct. T7 promoters and terminators are indicated by the arrows and crosses, respectively. **B.** tGFP assay principle: in the case of two interacting BMC-H, the GFP10 and 11 tags will come closer to each other. Reconstitution of a full fluorescent GFP will therefore be promoted in the presence of the GFP1-9 portion. Conversely, GFP reconstitution will be inefficient with non-interacting BMC-Hs.

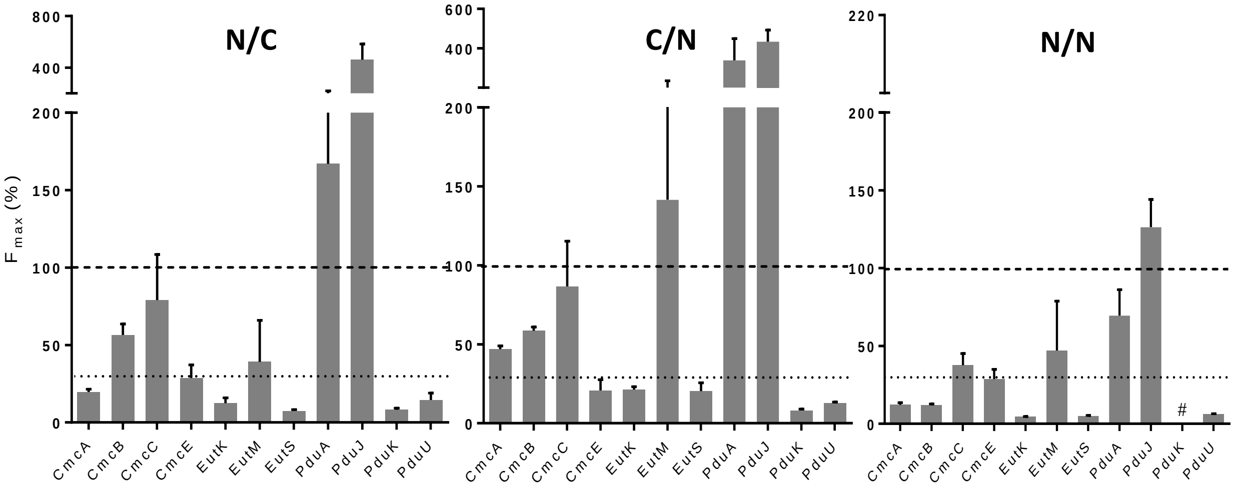

**Figure S4. Impact of GFP tag orientation scheme on tGFP signals.**

Fluorescence signals deriving from co-expression of Kpe BMC-H homo-pairs with GFP10/GFP11 tags attached following different configurations: N/C, C/N, or N/N orientations. The preparation of plasmid corresponding to the N/N PduK combination failed and could not be assayed (# symbol). Other experimental and data analysis details were as for Fig. 5.

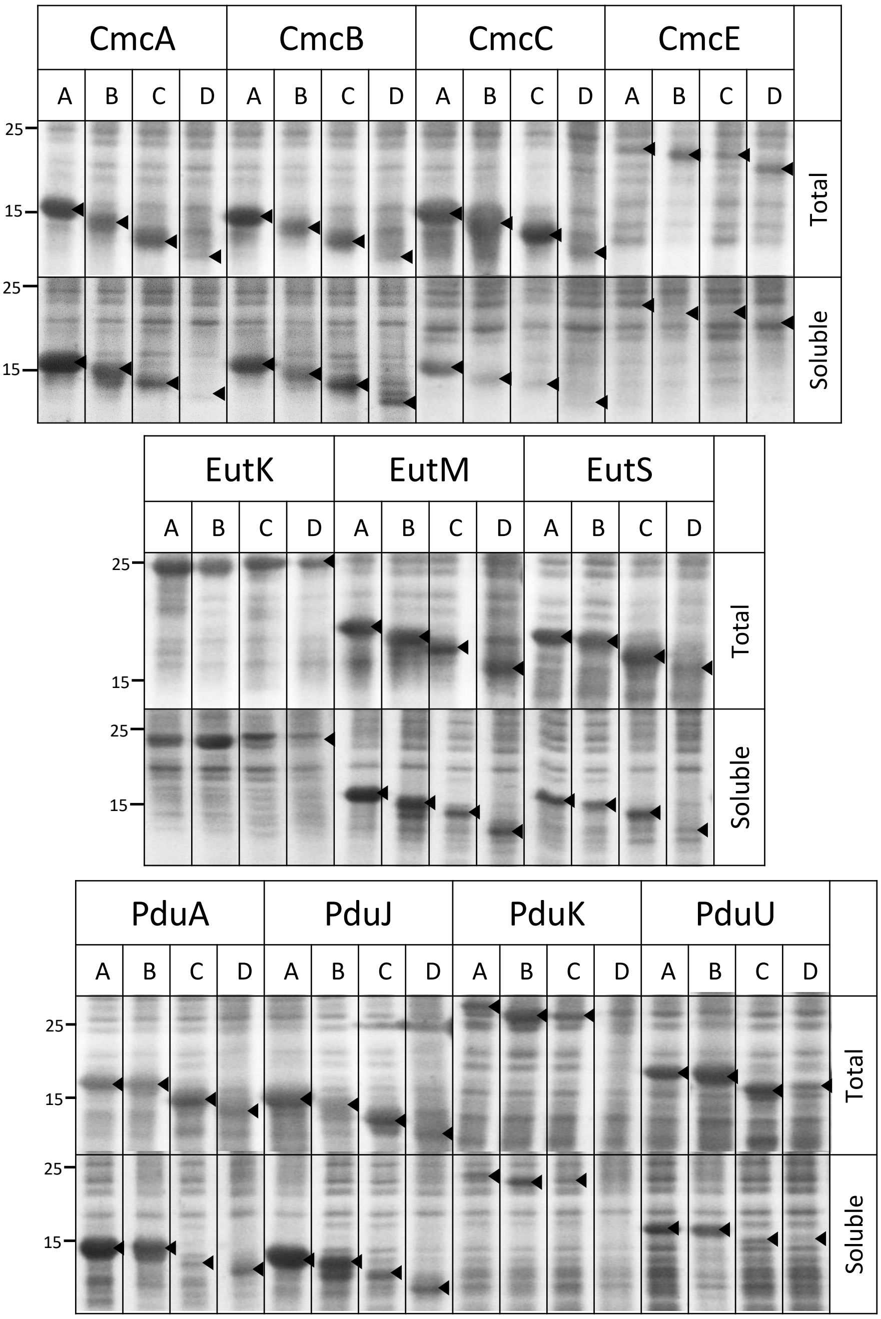

**Figure S5. Impact of GFP10/11 tag attribution and orientation on BMC-H expression and solubility.**

Individual BMC-H either tagged with GFP10 on C- (A) or N-terminus (B), or with GFP11 on the C- (C) or N-terminus (D) were over-expressed in BL21(DE3). After recovery of total cellular fractions, centrifugation permitted to prepare fractions corresponding to soluble contents. All fractions were analysed by SDS-PAGE and stained with Instant blue (Expedeon). The approximate migration of ladder components with indicated MW (kDa) are indicated on the left. Black arrows are given as an attempt to identify corresponding BMC-H bands. Theoretical MWs of BMC-H monomer constructs (kDa) are: CmcA 9.4; CmcB 9.6; CmcC 9.5; CmcE 13.6; EutK 17.0; EutM 9.8; EutS 11.6; PduA 9.8; PduJ 9.2; PduK 16.2; PduU 12.4. To these, it is necessary to add the contribution of linkers and tags, which differ as follows: for N-ter GFP10, 4.4; for C-ter GFP10, 4.6; for N-ter or C-ter GFP11, 4.7.

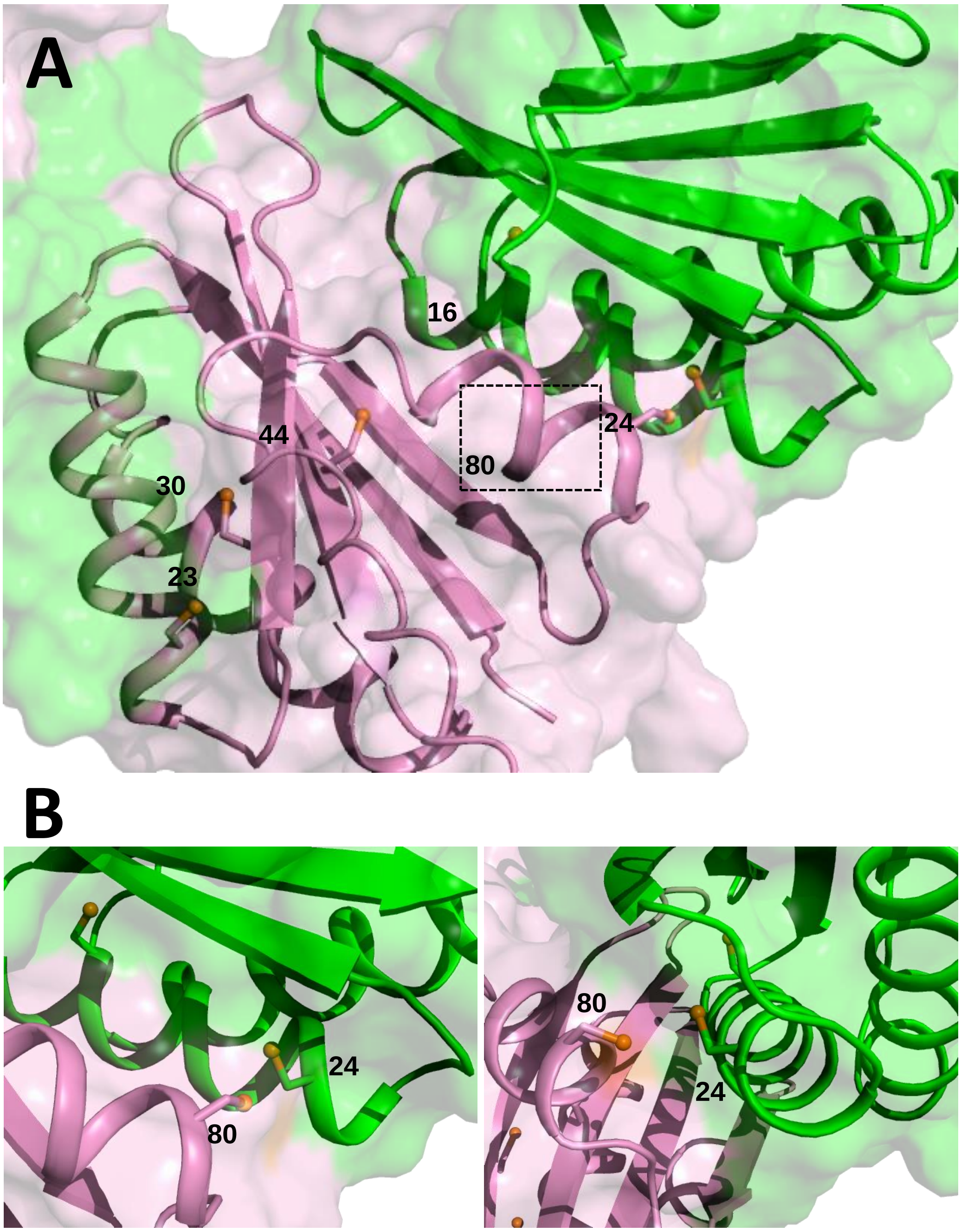

**Figure S6. Structural details predicted for GRM2 hetero-hexamers.**

***A,*** A potential disulfide bridge between residues Cys24 and Cys80 of CmcA (geen) and CmcE (violet), respectively, might stabilize contacts between monomers in hetero-hexameric associations. Depicted model corresponds to the AF2 prediction, but a similar result was obtained with ESMFold. Only two neighboring monomers are shown in cartoon representation to facilitate visualization. Cysteine sidechains are depicted as sticks with sulfur atoms as orange balls. Similar arrangement of cysteines is found when CmcA is replaced by CmcB or CmcC. ***B***, magnified views of the region around $CmcA^{24}/CmcE^{80}$.

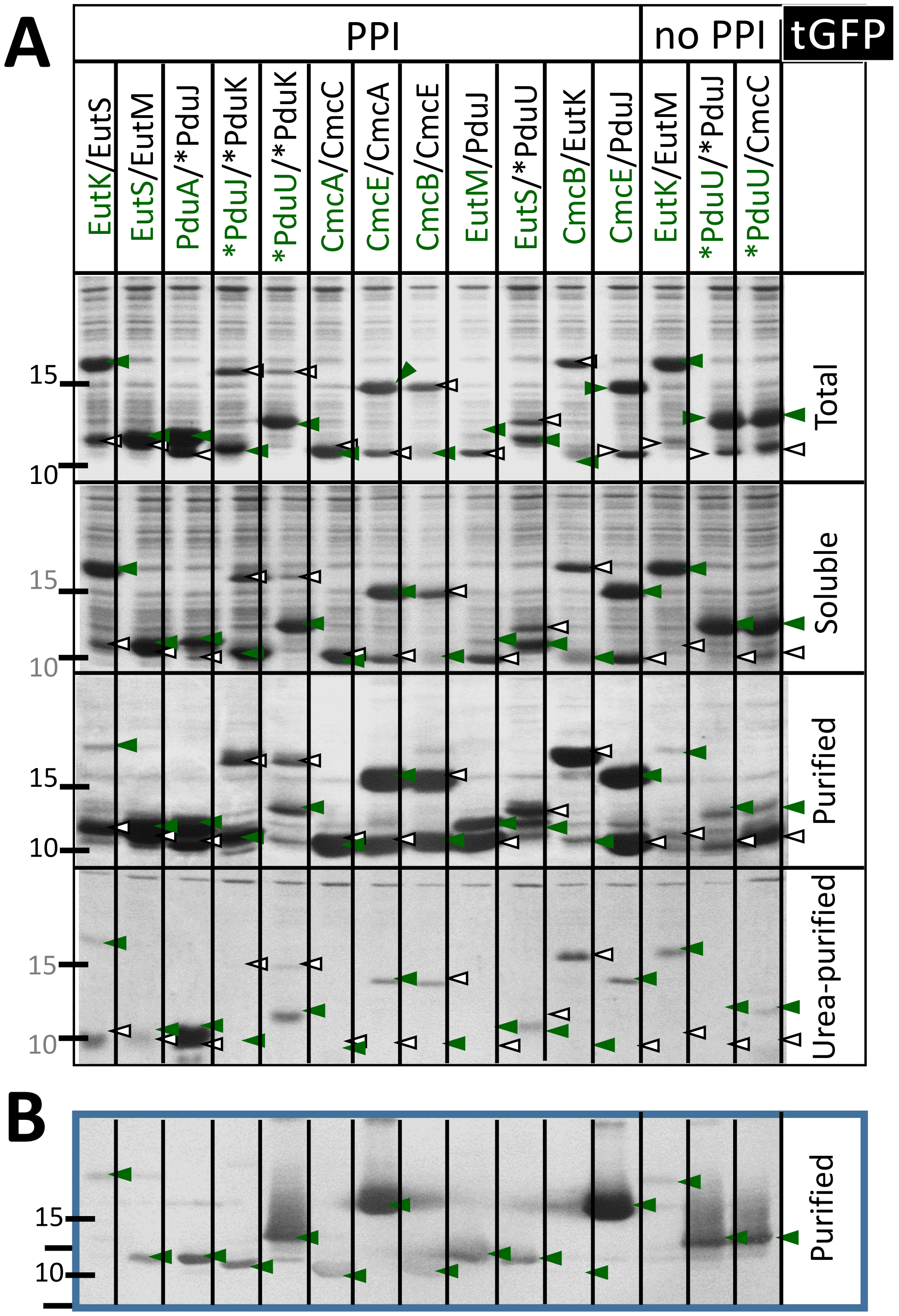

**Figure S7. Verification of *Kpe* BMC-H heteromerization.**
All details are as in Figure 6, with the exception that fractions were treated in the absence of β-mercaptoethanol.

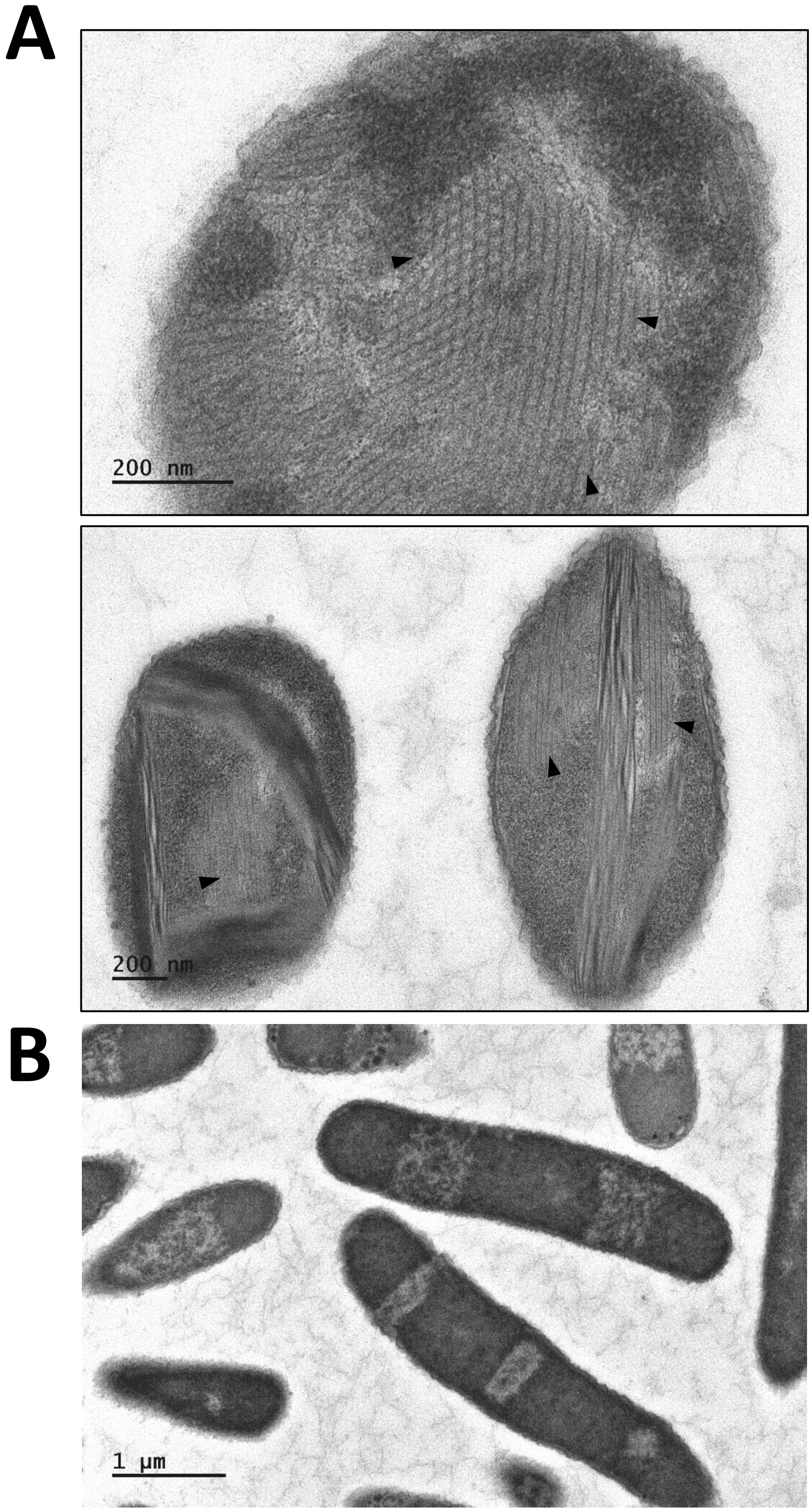

**Figure S8. BMC-H heteromers assembly.** TEM images showing contents of *E. coli* cells after over-expression of hetero-pairs combining PduJ-FLAG/PduA-His$_6$ (***A***) or CmcA-FLAG/CmcC-His6 (***B***). Regions displaying nanotubes are indicated by the arrows.

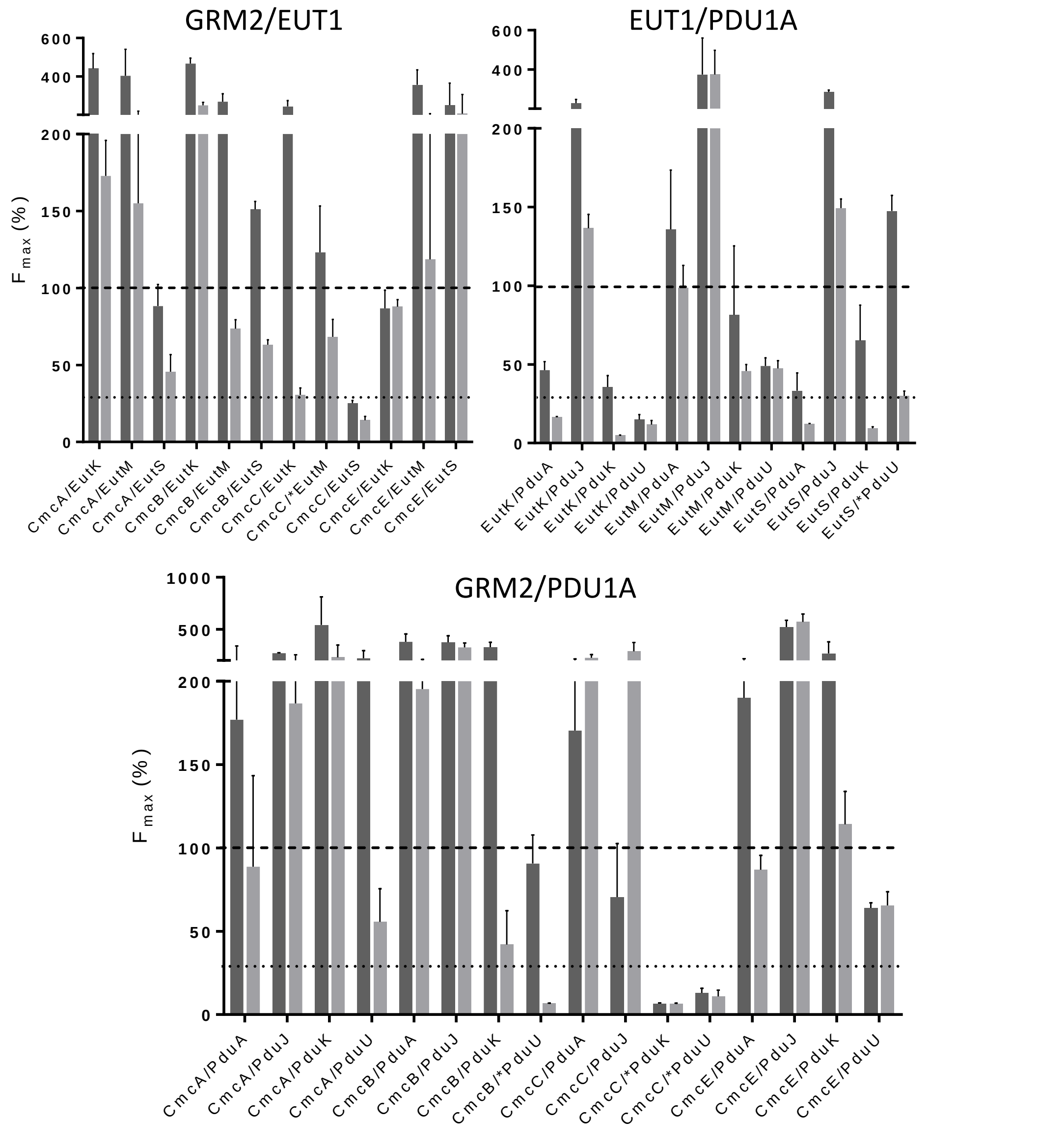

**Figure S9. Heteromeric association of BMC-H proteins arising from different BMC types.**

Please refer to figure 5 for details on data presentation.

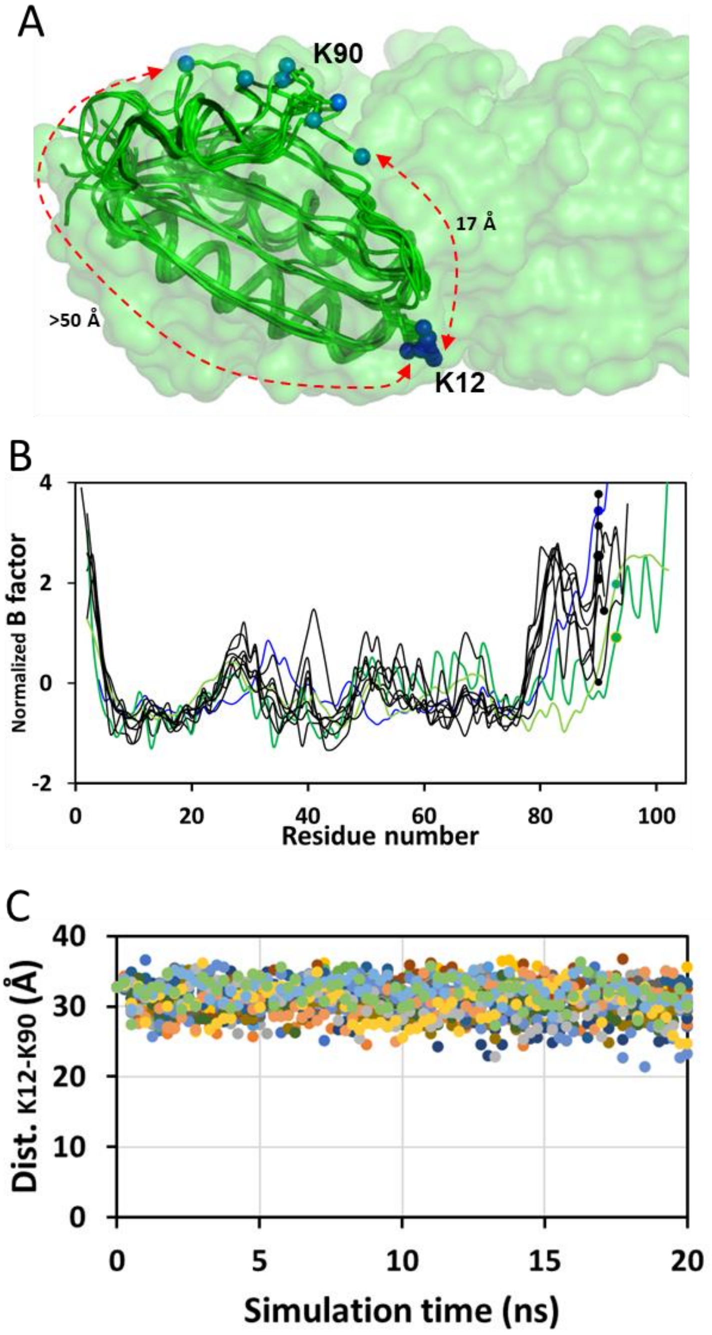

**Figure S10. Evaluation of the structural flexibility of the PduA K90 region.**
**A.** Comparison of the disposition of K12 and K90 side-chains in deposited 3D structures of PduA variants does not support big displacements towards the hexamer edge. C-terminal residues (including K90) are instead most often modelled towards the central pore. Wild-type PduA (3NGK) is shown in black trace, in green for all mutants (4PPD, 4QIE, 4QIF, 4GIG, 4RBT, 4RBU, 4RBV). Terminal K12 and K90 amine groups appear as spheres. **B.** Comparison of normalized crystallographic B temperature factors for Cα atoms in the different structures do not support extensive movements of residues preceding the PduA Lys90. The blue trace is for 3NGK, black for PduA mutants and green for two CcmK2 structures discussed by Trettel *et al*[41] (4OX7 and 21AB). Normalized B factor values seem indeed comparable between PduA K90 and the corresponding Arg93 of CcmK2 (dots, following the same color codes). This position precedes a region that is not highly mobile, according to MD simulations discussed in ref 41. Values were averaged over equivalent modeled residues in the hexamer. **C.** Molecular dynamic analysis does not support K90 extensive movements. Plotted are the evolution of K12/K90 inter-amine distances during 20ns run simulations of a PduA trihexameric assembly. Values never differed by more than 10 Å from the initial position, which is that from the crystal structure (3NGK). Shifts indicating approaches were due to movements towards the central pore. For each snapshot, 18 measurements are presented, corresponding to all the monomers from the 3NGK PduA tri-hexameric assembly studied before [57]. Note that inter-amine distances are measured in straight line, which is still incompatible with reaction in most instances.

**Table S1. Induction of *Kpe* BMC transcription in response to metabolite presence.**

| Substrates | Exp.[a] | SEM[b] | P-value | Exp.[a] | SEM[b] | P-value | Exp.[a] | SEM[b] | P-value |
|---|---|---|---|---|---|---|---|---|---|
| | *cmcA* | | | *cutC* | | | *cmcE* | | |
| w/o | 1.0 | 0.8<br>1.3 | / | 1.0 | 0.9<br>1.1 | / | 1.0 | 0.9<br>1.1 | / |
| EA | 1.0 | 0.4<br>2.3 | 0.966 | 0.4 | 0.2<br>0.9 | 0.335 | 0.2 | 0.1<br>0.3 | 0.057 |
| PD | 0.4 | 0.2<br>0.7 | 0.219 | 2.4 | 1.7<br>3.5 | 0.077 | 0.6 | 0.4<br>0.9 | 0.266 |
| CL | 476.9 | 422.2<br>538.7 | < 0.001 | 76.0 | 59,0<br>97.9 | < 0.001 | 22.9 | 18.5<br>28.4 | < 0.001 |
| EA + PD | 0.8 | 0.3<br>1.8 | 0.797 | 0.3 | 0.3<br>0.4 | 0.003 | 0.2 | 0.2<br>0.3 | 0.004 |
| EA + CL | 17.9 | 9.8<br>32.9 | 0.012 | 2.8 | 1.2<br>6.3 | 0.287 | 1.0 | 0.6<br>1.8 | 0.987 |
| PD + CL | 602.6 | 535.2<br>678.5 | < 0.001 | 67.0 | 54.6<br>82.3 | < 0.001 | 17.0 | 13.7<br>21.1 | < 0.001 |
| | *eutS* | | | *eutM* | | | *eutK* | | |
| w/o | 1.0 | 1,0<br>1,0 | / | 1.0 | 0.8<br>1.2 | / | 1.0 | 0.8<br>1.3 | / |
| EA | 1.1 | 0.8<br>1.4 | 0.831 | 3.8 | 3.2<br>4.4 | 0.006 | 0.4 | 0.3<br>0.5 | 0.083 |
| PD | 0.7 | 0.6<br>0.7 | 0.002 | 2.2 | 1.7<br>2.9 | 0.081 | 1.0 | 0.9<br>1,0 | 0.922 |
| CL | 0.7 | 0.6<br>0.8 | 0.054 | 17.1 | 15,0<br>19.5 | < 0.001 | 1.3 | 1.2<br>1.4 | 0.422 |
| EA + PD | 0.6 | 0.5<br>0.6 | 0.005 | 9.2 | 6.7<br>12.6 | 0.004 | 0.3 | 0.3<br>0.3 | 0.015 |
| EA + CL | 13.1 | 2.7<br>63.7 | 0.178 | 477.8 | 96.5<br>2366.9 | 0.019 | 5.7 | 1.5<br>22.1 | 0.273 |
| PD + CL | 0.7 | 0.6<br>0.7 | 0.016 | 22.7 | 20.1<br>25.7 | < 0.001 | 1.0 | 1,0<br>1.1 | 0.944 |
| | *pduA* | | | *pduJ* | | | *pduU* | | |
| w/o | 1.0 | 0.8<br>1.3 | / | 1.0 | 0.7<br>1.5 | / | 1.0 | 0.8<br>1.2 | / |
| EA | 0.1 | 0,0<br>0.1 | 0.027 | 0.3 | 0.1<br>0.9 | 0.344 | 0.9 | 0.3<br>2.7 | 0.929 |
| PD | 0.8 | 0.7<br>1,0 | 0.516 | 3.8 | 2.4<br>5.9 | 0.095 | 5.7 | 3,0<br>10.7 | 0.059 |
| CL | 5.1 | 4.3<br>6,0 | 0.005 | 36.2 | 28.8<br>45.6 | 0.002 | 26.7 | 24.4<br>29.2 | < 0.001 |
| EA + PD | 1.5 | 1,0<br>2.2 | 0.437 | 0.4 | 0.4<br>0.5 | 0.139 | 0.8 | 0.5<br>1.2 | 0.641 |
| EA + CL | 0.6 | 0.5<br>0.6 | 0.106 | 2.7 | 2,0<br>3.6 | 0.117 | 1.7 | 1.1<br>2.5 | 0.32 |
| PD + CL | 19.0 | 17.1<br>21,0 | < 0.001 | 78.7 | 63.9<br>96.9 | < 0.001 | 47.4 | 36.8<br>61.2 | < 0.001 |

[a] Expressions values are calculated from three independent replicates, and are given as fold changes relative to the condition without substrate.
[b] SEM values correspond to the expression lower and upper error bars.
Control measurements of housekeeping gene expression are compiled in Table S2.

**Table S2. Reproducibility of transcription level measurements of housekeeping genes.**

| Target genes | Substrates | Mean Cq[a] |
|---|---|---|
| *proC* | w/o | 28.38 |
| | | 27.64 |
| | | 27.63 |
| | EA | 26.16 |
| | | 25,00 |
| | | 23.86 |
| | PD | 26.29 |
| | | 25.88 |
| | | 26.37 |
| | CL | 24.67 |
| | | 24.25 |
| | | 24.18 |
| | EA + PD | 25.38 |
| | | 25.98 |
| | | 26.53 |
| | EA + CL | 24.55 |
| | | 24.57 |
| | | 25.32 |
| | PD + CL | 24.87 |
| | | 24.82 |
| | | 25.14 |

| Target genes | Substrates | Mean Cq[a] |
|---|---|---|
| *rpoD* | w/o | 27.24 |
| | | 26.60 |
| | | 26.53 |
| | EA | 24.66 |
| | | 23.81 |
| | | 23.19 |
| | PD | 28.26 |
| | | 27.29 |
| | | 27.76 |
| | CL | 24.59 |
| | | 24.29 |
| | | 24.46 |
| | EA + PD | 24.48 |
| | | 25.23 |
| | | 25.54 |
| | EA + CL | 24.45 |
| | | 24.99 |
| | | 24.63 |
| | PD + CL | 25.11 |
| | | 25.02 |
| | | 25,00 |

[a] mean quantification cycle, determined from three technical replicates.

**Table S3. Analysis of ESMFold and AF2 predictions for homo-hexamers**

| Protein | Hex[a] | pLDDT[b] | ic_PAE[c] | Interface[d] | | Core[e] | ΔE[f] |
|---|---|---|---|---|---|---|---|
| | | | | if_pLDDT | if_PAE | c_PAE | |
| **AlphaFold2** | | | | | | | |
| **CmcA** | YES | 95,8 | 1,7 | 96,6 | 1,4 | - | -63,5 |
| **CmcB** | YES | 95,6 | 1,8 | 95,2 | 1,6 | - | -58,6 |
| **CmcC** | YES | 94,9 | 1,9 | 95,6 | 1,5 | - | -54,9 |
| **EutM** | YES | 95,4 | 1,9 | 94,8 | 3,4 | - | -80,3 |
| **PduA** | YES | 93,4 | 2,3 | 95,6 | 3,1 | - | -62,9 |
| **PduJ** | YES | 94,5 | 2,0 | 95,1 | 1,6 | - | -61,1 |
| **CmcE** | YES | 72,8 | 7,4 | 90,4 | 2,8 | 1,9 | -54,4 |
| **EutK** | YES | 82,8 | 5,7 | 85,9 | 6,6 | 1,6 | -80,2 |
| **PduK** | YES | 67,7 | 7,7 | 73,1 | 9,5 | 3,0 | -51,3 |
| **EutS** | YES | 96,1 | 1,7 | 95,8 | 3,0 | 1,4 | -70,1 |
| **PduU** | YES | 92,6 | 2,5 | 95,6 | 1,6 | 1,8 | -77,2 |
| **BWI** | NO | 76,2 | 7,5 | - | - | - | - |
| **CcmK3** | YES | 96,2 | 1,8 | 96,3 | 1,5 | - | -72,0 |
| **RMM** | YES | 93,6 | 2,3 | 96,6 | 2,8 | - | -71,6 |
| **ESMFold** | | | | | | | |
| **CmcA** | YES | 80,1 | 2,9 | 86,4 | 1,9 | - | -54,3 |
| **CmcB** | YES | 78,9 | 3,1 | 84,0 | 2,0 | - | -61,7 |
| **CmcC** | YES | 81,9 | 2,6 | 87,4 | 1,7 | - | -53,8 |
| **EutM** | YES | 75,9 | 3,4 | 84,7 | 1,8 | - | -61,5 |
| **PduA** | YES | 76,8 | 3,4 | 80,6 | 2,7 | - | -53,4 |
| **PduJ** | YES | 80,9 | 2,9 | 85,7 | 2,0 | - | -54,7 |
| **CmcE** | YES | 59,2 | 8,5 | 71,6 | 7,8 | 4,2 | -14,2 |
| **EutK** | YES | 64,7 | 9,3 | 79,5 | 3,0 | 3,0 | -53,1 |
| **PduK** | NO | 47,1 | 12,3 | - | - | 9,3 | - |
| **EutS** | NO | 63,2 | 9,2 | - | - | 8,5 | - |
| **PduU** | YES | 62,8 | 5,2 | 64,6 | 7,3 | 4,5 | -61,3 |
| **BWI** | NO | 69,8 | 12,0 | - | - | - | - |
| **CcmK3** | NO | 59,3 | 7,3 | - | - | - | - |
| **RMM** | YES | 78,1 | 3,3 | 83,7 | 2,2 | - | -64,2 |

[a] Successful prediction of PF00936 hexameric associations is specified as YES/NO. [b] Global pLDDT. [c] interchain PAE, between pairs of residues belonging to different chains. [d] Values between pairs of interchain residues lying closer than 4 Å from each other. [e] values for interchain residues belonging to the BMC-H core, only calculated for non-canonical BMC-H. [f] Interaction energy (*ΔE*) computed using Rosetta InterfaceAnalyzer after structure relaxation, and averaged over all interfaces. Values of if_pLDDT, if_PAE and *ΔE* are reported only for combinations predicted as PF00936 hexamers.

**Table S4. Evaluation of inter-monomer interaction energies from experimental 3D structures**

| Name | RCSB | Origin | Commentary | ΔE[a] |
|---|---|---|---|---|
| BMC-H | 4QIV | *Aer. hydrophila* | | -59,0 |
| BMC-H | 4OLO | *Clos. bacterium* | apo form (Fe-S cluster) | -51,5 |
| BMC-H | 5DJB | *Hal. ochraceum* | | -65,0 |
| BMC-H | 6NLU | *Hal. ochraceum* | with circular permutation | -62,7 |
| CcmK1 | 3BN4 | *Syn. sp. PCC 6803* | | -77,5 |
| CcmK1 | 3DN9 | *Syn. sp. PCC 6803* | missing C-terminal residues | -69,6 |
| CcmK2 | 2A1B | *Syn. sp. PCC 6803* | | -66,9 |
| CcmK2 | 3CIM | *Syn. sp. PCC 6803* | missing C-terminal residues | -75,5 |
| CcmK2 | 3SSQ | *Ther. elongatus BP-1* | stacked hexamer | -86,7 |
| CcmK2 | 4OX7 | *Syn. elon. PCC 7942* | | -80,1 |
| CcmK2 | 3DNC | *Syn. sp. PCC 6803* | missing C-terminal residues | -75,5 |
| CcmK4 | 2A10 | *Syn. sp. PCC 6803* | | -82,6 |
| CcmK4 | 6SCR | *Syn. sp. PCC 6803* | | -80,7 |
| CcmK4 | 2A18 | *Syn. sp. PCC 6803* | | -80,1 |
| CcmK4 | 5VGU | *Hal. sp. PCC 7418* | | -62,8 |
| CcmK4 | 4OX6 | *Syn. elon. PCC 7942* | | -83,5 |
| CmcA | 7MGP | *E. coli* | K25A mutant | -65,1 |
| CmcB | 7MPW | *E. coli* | K25D mutant | -46,0 |
| CmcC | 7MPV | *E. coli* | K25A mutant | -57,2 |
| CsoS1 | 4OX8 | *Pro. marinus* | | -65,7 |
| CsoS1A | 2EWH | *Hal. neapolitanus* | | -69,5 |
| CsoS1A | 2G13 | *Hal. neapolitanus* | | -71,3 |
| CsoS1C | 3H8Y | *Hal. neapolitanus* | | -71,4 |
| CutN | 7MMX | *Strep. intermedius* | K27D mutant | -63,0 |
| CutR | 6XPI | *Strep. intermedius* | | -70,8 |
| CutR | 6XPK | *Strep. intermedius* | Screw-shaped hexamer | -35,1 |
| EutM | 3I6P | *E. coli* | | -72,3 |
| EutM | 3MPW | *E. coli* | | -66,3 |
| EutM | 3MPY | *E. coli* | | -73,1 |
| EutM | 4AXJ | *Clos. difficile* | | -63,7 |
| EutS | 3I96 | *E. coli* | dimer of trimers (ben hexamer) | -69,8 |
| EutS | 3IA0 | *E. coli* | G39V mutant, flat hexamer | -79,7 |
| PduA | 3NGK | *Sal. enterica* | | -56,0 |
| PduA | 4P7T | *Citr. freundii* | K26D mutant | -50,6 |
| PduA | 4P7V | *Citr. freundii* | K26D mutant | -56,6 |
| PduJ | 5D6V | *Sal. enterica* | Wild-type K25 manually reintroduced | -55,8 |
| PduU | 3CGI | *Sal. enterica* | | -76,4 |
| RMM | 5L38 | *Myc. Smegmatis* | | -69,1 |

[a] Averaged over all interfaces present in the 3D structure

**Table S5 –Assessment of *Kpe* BMC-H oligomeric state by SEC-HPLC.**

| | BMC-H(His6) | | |
|---|---|---|---|
| | BMC-H | MW[a] | SEC-MW[b] |
| EUT1 | EutK | 18.2 | 9/17 (1-M) |
| EUT1 | EutM [c] | 11.1 | 67 (2-M) |
| EUT1 | EutS [c] | 12.9 | 62 (2-M) |
| PDU1A | PduA | 10.8 | 58 (1-M) |
| PDU1A | PduJ | 10.3 | 58 (2-M)<br>9 (1-M) |
| PDU1A | *PduK [c] | 17.6 | 22 (2-M) |
| PDU1A | *PduU [c] | 13.8 | 71 (m) |
| GRM2 | CmcA | 10.6 | 67 (1-M) |
| GRM2 | CmcB | 10.8 | 72 (1-M) |
| GRM2 | CmcC | 10.6 | 68 (m)<br>21 (1-M) |
| GRM2 | CmcE [c] | 14.7 | 71 (m)<br>13 (2-M) |
| GRM2 | *CmcE [c] | 15.0 | 71 (m) |

| BMC-H(FLAG)/BMC-H(His6) | | | |
|---|---|---|---|
| BMC-H | MW[a] | SEC-MW[b] | |
| EutK/EutS [c] | 18.4/12.9 | 45 (2-M) | EUT1 |
| EutS/EutM [c] | 13.1/11.1 | 67 (2-M) | EUT1 |
| PduA/*PduJ [c] | 11.0/10.6 | 76 (1-M) | PDU1A |
| *PduJ/*PduK [c] | 10.8/17.6 | 110 (1-M) | PDU1A |
| *PduU/*PduK [c] | 14.0/17.6 | ND | PDU1A |
| CmcA/CmcC | 10.8/10.6 | 90 (1-M)<br>39 (2-M) | GRM2 |
| CmcE/CmcA | 14.9/10.6 | 153 (1-M)<br>34 (2-M) | GRM2 |
| CmcB/CmcE | 11.0/14.7 | 120 (1-M) | GRM2 |
| EutM/PduJ | 11.3/10.3 | 73 (1-M) | HYBRID |
| EutS/*PduU [c] | 13.1/13.8 | ND | HYBRID |
| CmcB/EutK | 11.0/18.2 | 12 (1-M) | HYBRID |
| CmcE/PduJ | 14.9/10.3 | 69 (1-M)<br>30 (2-M) | HYBRID |

[a]Theoretical molecular weight (MW) of monomer, in kDa. [b] MW estimated from peak elution volume, in kDa. Labels 1-M and 2-M are to indicate first or second major species in intensity, whereas m is for minor or faintly detected species; ND for nothing detected within the 5-500 kDa resolving range of the column. [c] These samples revealed peaks eluting as high-MW soluble species of highest intensity (1-M). Their retention time was between those of ferritin (440kDa) and dextran blue (2MDa), or even above the latter. Asterisks indicate tag attachment to the BMC-H N-terminus.

**Table S6. Analysis of ESMFold and AF2 predictions for hetero-hexamers combining monomers from the same BMC type.**

| BMC-H combination | Hex | Org.[a] | pLDDT | Ic_PAE | Interface | | Core | ΔE[b] | |
|---|---|---|---|---|---|---|---|---|---|
| | | | | | pLDDT | PAE | c_PAE | A/B | B/A |
| **AlphaFold2** | | | | | | | | | |
| **CmcA/CmcB** | YES | ABABAB | 94,4 | 2,0 | 94,8 | 3,8 | - | -67,4 | -63,7 |
| **CmcA/CmcC** | YES | ABABAB | 94,2 | 2,1 | 94,7 | 3,0 | - | -62,6 | -57,3 |
| **CmcA/CmcE** | YES | ABABAB | 81,2 | 5,5 | 85,8 | 8,7 | 4,6 | -65,1 | -60,0 |
| **CmcB/CmcC** | YES | ABABAB | 94 | 2,1 | 95,5 | 2,9 | - | -66,8 | -57,9 |
| **CmcB/CmcE** | YES | ABABAB | 81,2 | 5,4 | 85,6 | 8,7 | 4,4 | -72,3 | -63,3 |
| **CmcC/CmcE** | YES | ABABAB | 81,4 | 5,4 | 87 | 8,5 | 4,6 | -65,1 | -61,3 |
| **EutK/EutM** | YES | ABABAB | 88,2 | 5,8 | 91,4 | 11,5 | 4,8 | -65,2 | -89,0 |
| **EutK/EutS** | YES | AAABBB | 71,2 | 9,3 | 73,7 | 19,2 | 7,2 | - | - |
| **EutM/EutS** | YES | AAABBB | 83,4 | 5,4 | 74,9 | 12,4 | 5,2 | - | - |
| **PduA/PduJ** | YES | ABABAB | 93,5 | 2,2 | 94,9 | 3,9 | - | -63,0 | -63,1 |
| **PduA/PduK** | YES | ABABAB | 81,7 | 6,6 | 87 | 12,0 | 5,6 | -63,0 | -50,8 |
| **PduA/PduU** | YES | AAABBB | 79,4 | 6,6 | 78,4 | 15,8 | 6,3 | - | - |
| **PduJ/PduK** | YES | ABABAB | 82 | 6,5 | 85,9 | 12,9 | 5,6 | -83,0 | -46,7 |
| **PduJ/PduU** | YES | AAABBB | 79,1 | 6,8 | 77,5 | 16,6 | 6,5 | - | - |
| **PduK/PduU** | YES | ABABAB | 65,6 | 9,5 | 72,4 | 21,0 | 7,4 | -26,2 | -52,0 |
| **CcmK1/CcmK2** | YES | ABABAB | 95,4 | 2,0 | 96,6 | 3,5 | - | -83,4 | -82,4 |
| **ESMFold** | | | | | | | | | |
| **CmcA/CmcB** | YES | ABABAB | 80,2 | 2,9 | 85,0 | 2,1 | - | -69,7 | -74,2 |
| **CmcA/CmcC** | YES | ABABAB | 80,4 | 2,9 | 86,7 | 1,8 | - | -58,7 | -72,1 |
| **CmcA/CmcE** | YES | ABABAB | 67,6 | 5,0 | 81,0 | 2,6 | 3,3 | -53,9 | -64,3 |
| **CmcB/CmcC** | YES | ABABAB | 79,6 | 3,0 | 84,1 | 2,3 | - | -66,7 | -69,2 |
| **CmcB/CmcE** | YES | ABABAB | 68,6 | 4,9 | 81,3 | 2,6 | 3,4 | -58,5 | -63,9 |
| **CmcC/CmcE** | YES | ABABAB | 66,8 | 4,7 | 80,0 | 2,8 | 3,4 | -55,9 | -70,2 |
| **EutK/EutM** | YES | ABABAB | 70,9 | 7,2 | 80,8 | 2,8 | 3,3 | -76,7 | -81,4 |
| **EutK/EutS** | YES | AAABBB | 65,4 | 10,1 | 78,8 | 7,1 | 4,9 | - | - |
| **EutM/EutS** | NO | - | 72,0 | 8,5 | - | - | 8,0 | - | - |
| **PduA/PduJ** | YES | ABABAB | 80,4 | 2,9 | 85,7 | 2,0 | - | -63,6 | -69,5 |
| **PduA/PduK** | YES | ABABAB | 63,9 | 6,9 | 81,7 | 2,5 | 3,3 | -52,7 | -68,1 |
| **PduA/PduU** | YES | AAABBB | 67,0 | 6,4 | 71,5 | 14,2 | 5,7 | - | - |
| **PduJ/PduK** | YES | ABABAB | 64,5 | 6,9 | 82,8 | 2,4 | 3,2 | -44,4 | -61,9 |
| **PduJ/PduU** | YES | AAABBB | 68,6 | 5,8 | 75,7 | 9,8 | 5,1 | - | - |
| **PduK/PduU** | YES | AAABBB | 52,4 | 8,6 | 59,1 | 18,0 | 6,0 | - | - |
| **CcmK1/CcmK2** | YES | ABABAB | 71,2 | 4,3 | 80,9 | 2,5 | - | -84,3 | -81,0 |

[a] Organization mode of monomers in the top ranked predicted model. A and B are to identify the first and second monomer types. [b] Energies are averaged over the two different sets of 3 identical interfaces from the ABABAB hexamers. The last line corresponds to structural predictions for combined CcmK1 and CcmK2 from *Syn6803*. Other details are as in Table S3.

**Table S7A. Analysis of AF2 predictions for hetero-hexamers combining monomers from different BMC types.** Data were organized as in Table S6.

| BMC-H combination | AlphaFold2 | | | | | | | | |
|---|---|---|---|---|---|---|---|---|---|
| | Hex | Org. | pLDDT | ic_PAE | Interface | | Core | ΔE | |
| | | | | | pLDDT | PAE | c_PAE | A/B | B/A |
| CmcA/EutK | YES | ABABAB | 85,8 | 6,1 | 84,5 | 13,1 | 5,2 | -71,4 | -63,4 |
| CmcA/EutM | YES | ABABAB | 95,2 | 1,9 | 96,0 | 2,9 | - | -70,2 | -70,4 |
| CmcA/EutS | YES | AAABBB | 84,3 | 4,8 | 80,1 | 9,9 | 4,6 | - | - |
| CmcB/EutK | YES | ABABAB | 86,3 | 6,2 | 83,8 | 13,7 | 5,2 | -74,2 | -63,4 |
| CmcB/EutM | YES | ABABAB | 94,6 | 2,0 | 95,8 | 2,9 | - | -71,0 | -69,5 |
| CmcB/EutS | YES | AAABBB | 82,2 | 5,5 | 78,9 | 14,3 | 3,8 | - | - |
| CmcC/EutK | YES | ABABAB | 85 | 5,8 | 83,5 | 12,7 | 4,9 | -79,4 | -64,5 |
| CmcC/EutM | YES | ABABAB | 94,2 | 2,0 | 95,0 | 3,2 | - | -62,6 | -70,1 |
| CmcC/EutS | YES | AAABBB | 84,2 | 4,7 | 80,0 | 9,5 | 9,4 | - | - |
| CmcE/EutK | YES | ABABAB | 76,4 | 8,1 | 82,1 | 12,3 | 4,0 | -53,5 | -85,5 |
| CmcE/EutM | YES | ABABAB | 83,9 | 4,9 | 84,2 | 12,2 | 4,0 | -71,5 | -69,8 |
| CmcE/EutS | YES | AAABBB | 61,2 | 10,4 | 72,6 | 20,4 | 9,4 | - | - |
| CmcA/PduA | YES | ABABAB | 94,2 | 2,1 | 95,6 | 2,9 | - | -62,3 | -59,1 |
| CmcA/PduJ | YES | ABABAB | 94,8 | 2,0 | 96,1 | 2,8 | - | -68,3 | -62,5 |
| CmcA/PduK | YES | ABABAB | 82,4 | 6,4 | 86,8 | 12,2 | 5,5 | -75,5 | -49,5 |
| CmcA/PduU | YES | AAABBB | 80,4 | 6,3 | 85,4 | 13,3 | 5,9 | - | - |
| CmcB/PduA | YES | ABABAB | 93,8 | 2,2 | 95,7 | 3,4 | - | -67,2 | -65,1 |
| CmcB/PduJ | YES | ABABAB | 95,2 | 1,9 | 96,6 | 2,7 | - | -67,3 | -64,2 |
| CmcB/PduK | YES | ABABAB | 82,1 | 6,5 | 86,1 | 12,3 | 5,5 | -71,1 | -56,1 |
| CmcB/PduU | YES | AAABBB | 81,4 | 5,9 | 75,1 | 12,3 | 5,2 | - | - |
| CmcC/PduA | YES | ABABAB | 93,2 | 2,2 | 93,9 | 3,1 | - | -59,2 | -59,5 |
| CmcC/PduJ | YES | ABABAB | 94,4 | 2,0 | 95,1 | 3,5 | - | -59,2 | -63,6 |
| CmcC/PduK | YES | ABABAB | 82,1 | 6,5 | 85,8 | 12,3 | 5,5 | -62,2 | -48,7 |
| CmcC/PduU | YES | AAABBB | 79,2 | 6,7 | 73,9 | 15,6 | 5,5 | - | - |
| CmcE/PduA | YES | ABABAB | 81,3 | 5,5 | 80,5 | 14,1 | 4,5 | -51,3 | -61,0 |
| CmcE/PduJ | YES | ABABAB | 81,4 | 5,4 | 79,4 | 13,4 | 4,5 | -56,4 | -59,2 |
| CmcE/PduK | YES | ABABAB | 72,2 | 8,1 | 79,1 | 11,3 | 4,0 | -72,1 | -46,6 |
| CmcE/PduU | YES | AAABBB | 62,3 | 10,1 | 65,1 | 18,9 | 8,6 | - | - |
| EutK/PduA | YES | ABABAB | 86,1 | 6,3 | 85,8 | 12,3 | 5,3 | -61,7 | -77,6 |
| EutK/PduJ | YES | ABABAB | 87,7 | 4,6 | 87,4 | 10,1 | 3,9 | -66,3 | -78,3 |
| EutK/PduK | YES | ABABAB | 74,2 | 8,3 | 84,8 | 7,2 | 2,7 | -68,0 | -47,9 |
| EutK/PduU | YES | AAABBB | 68,8 | 10,0 | 57,5 | 18,2 | 7,9 | - | - |
| EutM/PduA | YES | ABABAB | 93,5 | 2,3 | 93,9 | 3,4 | - | -70,0 | -65,5 |
| EutM/PduJ | YES | ABABAB | 94,6 | 2,0 | 95,2 | 3,5 | - | -69,9 | -63,4 |
| EutM/PduK | YES | ABABAB | 80,9 | 6,5 | 82,6 | 13,7 | 5,3 | -82,0 | -50,5 |
| EutM/PduU | YES | AAABBB | 77,6 | 8,0 | 67,7 | 18,3 | 7,4 | - | - |
| EutS/PduA | YES | AAABBB | 81,4 | 5,7 | 76,6 | 11,9 | 5,3 | - | - |
| EutS/PduJ | YES | AAABBB | 80,2 | 6,5 | 74,8 | 15,5 | 6,2 | - | - |
| EutS/PduK | YES | ABABAB | 65,8 | 9,4 | 70,8 | 21,8 | 7,6 | -51,9 | -24,2 |
| EutS/PduU | YES | ABABAB | 94,1 | 2,2 | 93,6 | 5,5 | 1,7 | - | - |
| CcmK1/CcmK2 | YES | ABABAB | 95,4 | 2,0 | 96,6 | 3,5 | - | -83,4 | -82,4 |

**Table S7B. Analysis of ESMFold predictions for hetero-hexamers combining monomers from different BMC types.** Please refer to Table S6 for data organization details.

| BMC-H combination | ESMFold | | | | | | | | |
|---|---|---|---|---|---|---|---|---|---|
| | Hex | Org. | pLDDT | ic_PAE | Interface | | Core | ΔE | |
| | | | | | pLDDT | PAE | c_PAE | A/B | B/A |
| CmcA/EutK | YES | ABABAB | 70,8 | 7,2 | 82,6 | 2,6 | 3,1 | -62,3 | -70,0 |
| CmcA/EutM | YES | ABABAB | 80,3 | 3,0 | 87,42 | 1,74 | - | -71,6 | -74,7 |
| CmcA/EutS | NO | - | 74,8 | 8,5 | - | - | 8,1 | - | - |
| CmcB/EutK | YES | ABABAB | 69,6 | 7,3 | 81,8 | 2,7 | 3,3 | -64,5 | -70,4 |
| CmcB/EutM | YES | ABABAB | 78,5 | 3,2 | 85,22 | 1,96 | - | -71,5 | -81,8 |
| CmcB/EutS | NO | - | 73,7 | 8,5 | - | - | 8,1 | - | - |
| CmcC/EutK | YES | ABABAB | 71,0 | 7,3 | 81,8 | 2,5 | 3,0 | -56,0 | -69,9 |
| CmcC/EutM | YES | ABABAB | 79,9 | 2,9 | 86,55 | 1,86 | - | -71,1 | -61,6 |
| CmcC/EutS | NO | - | 74,0 | 8,4 | - | - | 8,0 | - | - |
| CmcE/EutK | YES | ABABAB | 64,0 | 8,7 | 78,6 | 3,5 | 3,4 | -83,6 | -49,4 |
| CmcE/EutM | YES | ABABAB | 65,0 | 6,0 | 77,0 | 3,6 | 3,7 | -71,9 | -76,2 |
| CmcE/EutS | YES | AAABBB | 57,0 | 8,2 | 75,4 | 15,4 | 6,9 | - | - |
| CmcA/PduA | YES | ABABAB | 78,7 | 3,1 | 83,3 | 2,26 | - | -63,6 | -78,7 |
| CmcA/PduJ | YES | ABABAB | 81,6 | 2,8 | 85,93 | 1,99 | - | -61,1 | -68,6 |
| CmcA/PduK | YES | ABABAB | 63,6 | 7,0 | 82,1 | 2,4 | 3,5 | -50,1 | -63,9 |
| CmcA/PduU | NO | - | 68,7 | 6,6 | - | - | 5,9 | - | - |
| CmcB/PduA | YES | ABABAB | 78,2 | 3,2 | 83,95 | 2,15 | - | -68,0 | -71,5 |
| CmcB/PduJ | YES | ABABAB | 80,3 | 3,0 | 86,11 | 1,9 | - | -60,3 | -71,0 |
| CmcB/PduK | YES | ABABAB | 63,8 | 5,9 | 79,5 | 3,0 | 3,5 | -53,6 | -59,7 |
| CmcB/PduU | NO | - | 68,2 | 6,5 | - | - | 5,8 | - | - |
| CmcC/PduA | YES | ABABAB | 80,2 | 2,9 | 85,37 | 1,93 | - | -65,1 | -65,9 |
| CmcC/PduJ | YES | ABABAB | 81,8 | 2,7 | 86,59 | 1,85 | - | -59,7 | -64,0 |
| CmcC/PduK | YES | ABABAB | 64,2 | 6,9 | 81,9 | 2,5 | 3,2 | -48,5 | -54,7 |
| CmcC/PduU | YES | AAABBB | 68,8 | 6,6 | 76,42 | 13,16 | 5,9 | - | - |
| CmcE/PduA | YES | ABABAB | 65,8 | 5,9 | 79,9 | 2,8 | 3,4 | -60,8 | -52,1 |
| CmcE/PduJ | YES | ABABAB | 67,1 | 5,4 | 78,5 | 3,3 | 3,3 | -72,2 | -56,2 |
| CmcE/PduK | YES | ABABAB | 59,5 | 8,4 | 75,9 | 4,9 | 3,7 | -34,0 | -37,4 |
| CmcE/PduU | YES | AAABBB | 53,6 | 9,2 | 68,5 | 15,7 | 7,0 | - | - |
| EutK/PduA | YES | ABABAB | 68,4 | 7,2 | 77,9 | 3,3 | 3,5 | -70,2 | -67,5 |
| EutK/PduJ | YES | ABABAB | 70,3 | 7,1 | 80,2 | 2,9 | 3,2 | -66,8 | -66,0 |
| EutK/PduK | YES | ABABAB | 59,8 | 9,3 | 77,2 | 4,0 | 3,4 | -56,3 | -79,3 |
| EutK/PduU | YES | AAABBB | 63,2 | 10,0 | 74,3 | 7,0 | 4,5 | - | - |
| EutM/PduA | YES | ABABAB | 78,4 | 3,2 | 84,41 | 2,23 | - | -63,8 | -71,4 |
| EutM/PduJ | YES | ABABAB | 79,7 | 3,0 | 85,38 | 2,04 | - | -61,0 | -72,5 |
| EutM/PduK | YES | ABABAB | 64,1 | 7,1 | 81,3 | 2,7 | 3,4 | -69,2 | -65,2 |
| EutM/PduU | YES | AAABBB | 65,5 | 6,5 | 68,62 | 14,22 | 5,8 | - | - |
| EutS/PduA | NO | - | 72,0 | 7,2 | - | - | 6,8 | - | - |
| EutS/PduJ | NO | - | 72,0 | 8,3 | - | - | 7,9 | - | - |
| EutS/PduK | YES | AAABBB | 55,3 | 8,5 | 75,6 | 13,4 | 7,1 | - | - |
| EutS/PduU | YES | ABBAAB | 68,4 | 4,9 | 77,97 | 3,66 | 3,4 | - | - |
| CcmK1/CcmK2 | YES | ABABAB | 71,2 | 4,3 | 80,9 | 2,5 | - | -84,3 | -81,0 |

**Table S8.** Primers for RT-qPCR experiments

| Primer name | Primer sequence (5'- 3') | Use | Amplicon size (bp) |
|---|---|---|---|
| qPCR-eutS-Fw<br>qPCR-eutS-Rv | TCACACTGGCGCATCTGATT<br>CCCGGTGTCAGCGTCATAAT | Quantification of *eutS* expression by RT-qPCR | 100 |
| qPCR-eutM-Fw<br>qPCR-eutM-Rv | CGTATCGGTGAGCTGGTCTC<br>GCTATCGCCCTTGAAGCTGA | Quantification of *eutM* expression by RT-qPCR | 90 |
| qPCR-eutK-Fw<br>qPCR-eutK-Rv | TCCGGAAGAGGATACCCAGT<br>TAACGCTTCCGATGATGCCG | Quantification of *eutK* expression by RT-qPCR | 94 |
| qPCR-cmcA-Fw<br>qPCR-cmcA-Rv | GATGTGTAAAGCCGCCAACG<br>GACGTCGCCTTTCACCATCA | Quantification of *cmcA* expression by RT-qPCR | 85 |
| qPCR-cutC-Fw<br>qPCR-cutC-Rv | TTGACGGCTATCCGATGCTC<br>AACATGGCGGAGAGTTCGTT | Quantification of *cutC* expression by RT-qPCR | 82 |
| qPCR-cmcE-Fw<br>qPCR-cmcE-Rv | CATCCACACCGCCATTGAAC<br>CTTCAACCACACAGCGCTCC | Quantification of *cmcE* expression by RT-qPCR | 98 |
| qPCR-pduA-Fw<br>qPCR-pduA-Rv | AGGCTTAACCGCAGCCATAG<br>AACCGATCCTTTCGTAGCCC | Quantification of *pdA* expression by RT-qPCR | 83 |
| qPCR-pduJ-Fw<br>qPCR-pduJ-Rv | TTGAAGCCGCTGATGCAATG<br>GCGGACCATCACGGTAATCA | Quantification of *pduJ* expression by RT-qPCR | 92 |
| qPCR-pduU-Fw<br>qPCR-pduU-Rv | GCATTCTCACCATTACCCCCA<br>AAGCGATCGAGAAAGCCGAT | Quantification of *pduU* expression by RT-qPCR | 94 |
| qPCR-proC-Fw<br>qPCR-proC-Rv | GATTGCCGATATCGTCTTCG<br>GAGACCACCAGCGACTCTTT | Quantification of *proC* expression by RT-qPCR | 99 |
| qPCR-recA-Fw<br>qPCR-recA-Rv | TTAAACAGGCCGAATTCCAG<br>CCGCTTTCTCAATCAGCTTC | Quantification of *recA* expression by RT-qPCR | 99 |
